# Internal sugar allocation in response to a shade signal is regulated by concerted action of auxin and sucrose

**DOI:** 10.64898/2026.05.04.722612

**Authors:** Sandi Paulišić, Blanca Jazmin Reyes-Hernández, Yetkin Çaka Ince, Jade Heinel, Christian Fankhauser, Johanna Krahmer

## Abstract

Plants detect proximity to neighboring vegetation by light spectral signatures which are sensed by phytochrome photoreceptors. As a response, many species grow taller to out-compete their neighbors. In seedlings, the rapid elongation of hypocotyls requires enhanced supply of carbon resources from the cotyledons, transported as sucrose. The mechanisms of how phytochrome signaling regulates carbon allocation are unknown. We show that sucrose biosynthesis, particularly the step catalyzed by sucrose phosphate synthase (SPS), is a key determinant of hypocotyl elongation. Moreover, we show that auxin directs resource allocation to the elongating hypocotyl. Increased sucrose availability enhances elongation only when auxin levels are sufficient, highlighting the interdependence between sugar supply and auxin. In contrast, our data reveal that reduced sucrose availability does not impair neighbor-proximity-induced auxin synthesis and signaling in hypocotyls and cotyledons. Our findings shed light on the regulation of carbon allocation — a process which is poorly understood, despite its importance for crop yield.

## Introduction

Proximity to vegetation is a potential risk of future canopy shade for plants, resulting in responses preempting reduced access to light. Many species respond to neighbor cues by growth and architectural changes which serve to over-top their neighbors – referred to as ‘neighbor detection’. Such changes include elongation of hypocotyls, petioles and stems (Casal, 2013; Fiorucci & Fankhauser, 2017; Ballaré & Pierik, 2017), higher leaf elevation angles (Michaud *et al*., 2017; Pantazopoulou *et al*., 2017) and enhanced phototropism (Goyal *et al*., 2016).

Plants sense their neighbors via spectral signatures: Plant tissue absorbs most blue and red (R) light but very little far red (FR) light. Therefore, neighbors reflect mainly FR (700nm to 800nm), which increases the relative amount of FR in the ambient light environment. Plant density therefore determines the ratio of R to FR light (R/FR) with a lower R/FR indicating the presence of neighbors. R/FR is sensed by phytochrome (phy) photoreceptors. phyB is the main photoreceptor responsible for the neighbor detection response, with contributions from other phytochromes. phyB is activated by R light and inactivated by FR light, which explains its ability to sense R/FR (Casal, 2013; Fiorucci & Fankhauser, 2017; Ballaré & Pierik, 2017).

The molecular signaling pathways regulated by the activation status of phyB are well characterized, especially in the hypocotyl elongation response of seedlings. Activated phyB suppresses PHYTOCHROME INTERACING FACTORs (PIFs) which are transcription factors – especially PIF7 is important for neighbor detection signaling (Li *et al*., 2012). Reduced phyB activation therefore de-represses PIFs and allows them to activate their targets. Among the genes activated by PIFs are the *YUCCA* genes which code for enzymes in the canonical biosynthetic pathway of the growth hormone auxin (Fiorucci & Fankhauser, 2017). Therefore, at low R/FR, more auxin is produced, mainly in cotyledons of *Arabidopsis* and *Brassica rapa* seedlings. Auxin is then transported to the hypocotyl of seedlings where it promotes elongation (Procko *et al*., 2014; Fiorucci & Fankhauser, 2017). Other mechanisms contribute in addition to this major pathway, such as local production of auxin in the hypocotyl (Zheng *et al*., 2016).

In contrast to the signaling pathways, the knowledge of the actual growth processes in hypocotyls is quite limited. Cell wall synthesis and modification, as well as re-organization of microtubules for example are critical, but their connections to neighbor detection signaling pathways are not well established (Yu *et al*., 2015; Ince *et al*., 2022; Sénéchal *et al*., 2024). Likewise, our knowledge of how elongation growth is fueled in low R/FR is limited (Krahmer & Fankhauser, 2024). During the low R/FR-induced elongation response of *Brassica rapa* hypocotyls and mustard internodes, more dry biomass is allocated to the elongating organs (Casal *et al*., 1995; Yanovsky *et al*., 1995; de Wit *et al*., 2018). Therefore, the observed hypocotyl elongation is not merely achieved through increased water or by a decrease of diameter to length ratio.

Moreover, in the case of mustard internode and *Brassica rapa* hypocotyl elongation, increased resource allocation to stem / hypocotyl is at the expense of carbon retained in leaves / cotyledons, indicating that the mustard stem and *Brassica rapa* hypocotyl are axial sinks, relying on import of carbon from leaves or cotyledons (Casal *et al*., 1995; Yanovsky *et al*., 1995; de Wit *et al*., 2018). Interestingly, in the case of end of day FR treatment of mustard plants, carbon export is only increased from leaves exposed to the FR pulse (Casal *et al*., 1995). Therefore, light signaling in the leaf regulates carbon allocation changes, which do not exclusively depend on increased carbon demand in elongating stems.

In many species, including *Arabidopsis thaliana* and *Brassica rapa*, carbon transport from source to sink organs takes place in the form of sucrose in the phloem sieve elements inside the vasculature (Braun, 2022). Arabidopsis mutants which are impaired in different aspects of sucrose transport – *suc2-4* and *sweet11/12* elongate less or not at all in response to low R/FR, suggesting that enhanced sucrose transport from cotyledons to hypocotyl is required for elongation (de Wit *et al*., 2018). By contrast, a mutant with constitutively high sugar levels, *pgm1-1*, has longer hypocotyls in high and low R/FR, indicating that high sugar abundance can cause hypocotyl elongation in the absence of a neighbor detection light signal (de Wit *et al*., 2018). In mustard leaves, the activity of the sucrose biosynthetic enzyme sucrose-phosphate synthase (SPS) is increased by an end of day FR pulse (Yanovsky *et al*., 1995). SPS activity is also enhanced in radish leaves by long-term exposure to low R/FR light (Keiller & Smith, 1989). While SPS catalyzes the synthesis of sucrose, invertase catalyzes sucrose breakdown into glucose and fructose – a process which can enhance unloading of sucrose from the phloem (Milne *et al*., 2018). Higher invertase activity was detected in petioles of radish plants grown in low R/FR, indicating that sucrose dynamics is affected by phytochrome deactivation not only in source but also in an elongating sink organ (Keiller & Smith, 1989).

In addition to the importance of sugar for carbon supply, hexoses and sucrose can also influence neighbor detection signaling. The *pgm pif7* mutant fails to elongate in high and low R/FR while sucrose levels are as high as in the *pgm* mutant, indicating that excess sugar can only cause elongation in the presence of an intact PIF dependent signaling pathway (de Wit *et al*., 2018). Furthermore, exogenous sucrose treatment induces hypocotyl elongation by boosting auxin biosynthesis via PIFs (Stewart *et al*., 2011; Lilley *et al*., 2012; Sairanen *et al*., 2012). The transcription factor *RSS1*, which inhibits hypocotyl elongation and is in turn inhibited by glucose, has been proposed as a link between glucose and auxin (Singh *et al*., 2017).

The molecular mechanisms by which phytochrome signaling controls resource allocation are unknown. Moreover, our mechanistic knowledge on the interplay between sugar, auxin and phytochrome signaling remains limited. In this paper, we focus on sucrose because enhanced carbon allocation to hypocotyls is essential for their elongation, and carbon is transported through the plant primarily as sucrose in the phloem. We firstly determine, which metabolic enzymes are required in the source tissue to produce sucrose to fuel hypocotyl elongation. Secondly, we investigated whether carbon allocation changes to the hypocotyl are mediated by auxin. Thirdly we determine the relationship between auxin and sucrose in this response.

## Materials and methods

### Plant material and growth conditions

*Arabidopsis thaliana* WT and mutants (all in the Col-*0* background) and WT plants of the *Brassica rapa* variety R-o-18 were used in this study. All *sps* mutants and combinations were kindly provided by Prof Javier Pozueta-Romero, Málaga University, Spain (Bahaji *et al*., 2015), the *f2kp* mutant (SALK_016314) (McCormick & Kruger, 2015) by Prof Tom Hamborg Nielsen, University of Copenhagen. Moreover, we used *cyfbp* (SALK_064456) (Cho & Yoo, 2011), *pgm1-1* (‘*pgm*’) (Caspar *et al*., 1985), *sav3-2* (*‘sav3’*) (Tao *et al*., 2008)*, suc2-4 (‘suc2’)* (Srivastava *et al*., 2008)*, FRO6:XVE:YUC3* (Chen *et al*., 2014) and pPDF1:DII-VENUS-N7-2A-TagBFP-sv40 (‘qDIIV’) (Galvan-Ampudia *et al*., 2018; Boccaccini *et al*., 2020). The *pgm sav3* mutant was generated by crossing *pgm1-1* with *sav3-2*, *pgm spsA1/C* by crossing *pgm1-1* with *spsA1/C*, the qDIIV in the *spsA1/C* background by crossing pPDF1:DII-VENUS-N7-2A-TagBFP-sv40 with *spsA1/C*.

In most experiments, light settings for high and low R/FR and LB were as used previously (Ince *et al*., 2022) (Table 1, Fig. S1A-E), using cool white fluorescent tubes and FR LEDs (peak emission 745nm, in GroLED module) in Percival plant growth chambers (Fig. S1A,B). For qDIIV (Fig. 4B,C), sugar measurements (Fig. 2A,C), DW and anthrone assay experiments (Fig. 3B-E, S2A-E), a Polyklima cabinet with cool white LEDs was used (Table 1, Fig. S1D,E). Light settings in the Polyklima cabinet were carefully adjusted such that PAR, elongation response and expected phytochrome activation state were equivalent to the experiments done in Percivals. The phytochrome activation state (P_fr_/P_tot_) was calculated by an in-house matlab script using oat phyA transition rate spectra (Mancinelli, 1994) which was kindly provided in digital format by Prof Christian Fleck, Wageningen University and Research.

**Figure 1:**
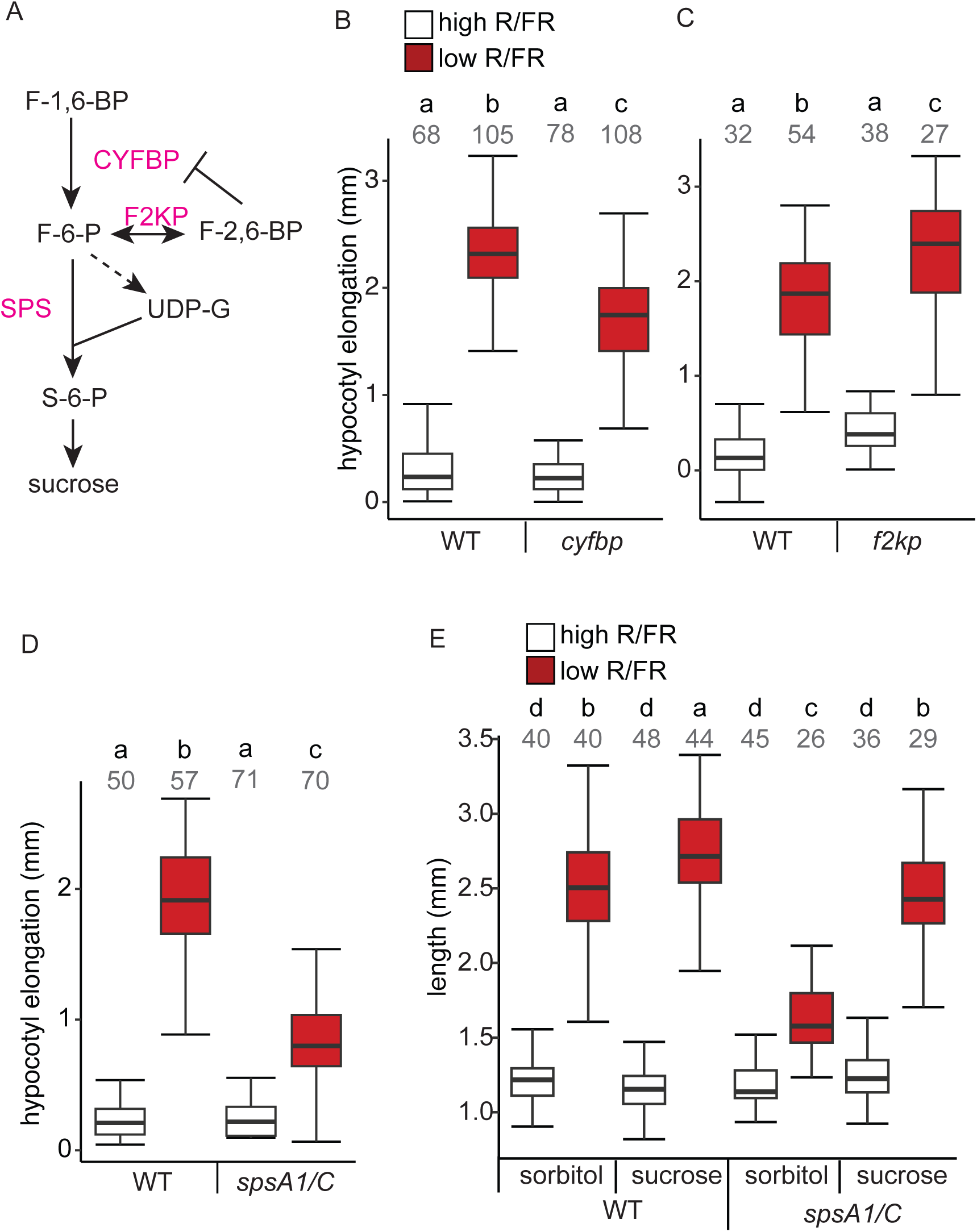
Sucrose synthesis capacity is neccessary hypocotyl elongation in the neighbor detection response. A) Overview of the sucrose biosynthetic pathway and relevant enzymes. F-6-P: fructose-6-phosphate, F-1,6-BP: fructose-1,6-bisphosphate, F-2,6-BP: fructose-2,6-bisphosphate, UDP-G: UDP-glucose, S-6-P: sucrose-6-phosphate. Enzymes are highlighted in magenta. B-D) Hypocotyl elongation between 5d and 8d in seedlings of *cyfbp* (B), *f2kp* (C) and *spsA1/C* (D), in high R/FR (white) or low R/FR (dark red). E) Hypocotyl length of 8-day old seedlings grown in high R/FR for 5 days followed by 3 days in either high or low R/FR and application of either 90mM sucrose or 90mM sorbitol to cotyledons. Grey numbers in (B-E) indicate the number of hypocotyls analyzed. Lower-case letters above samples in (B-E): different letters indicate significant difference between samples, ANOVA followed by Tukey-HSD test, P<0.05.

**Figure 2:**
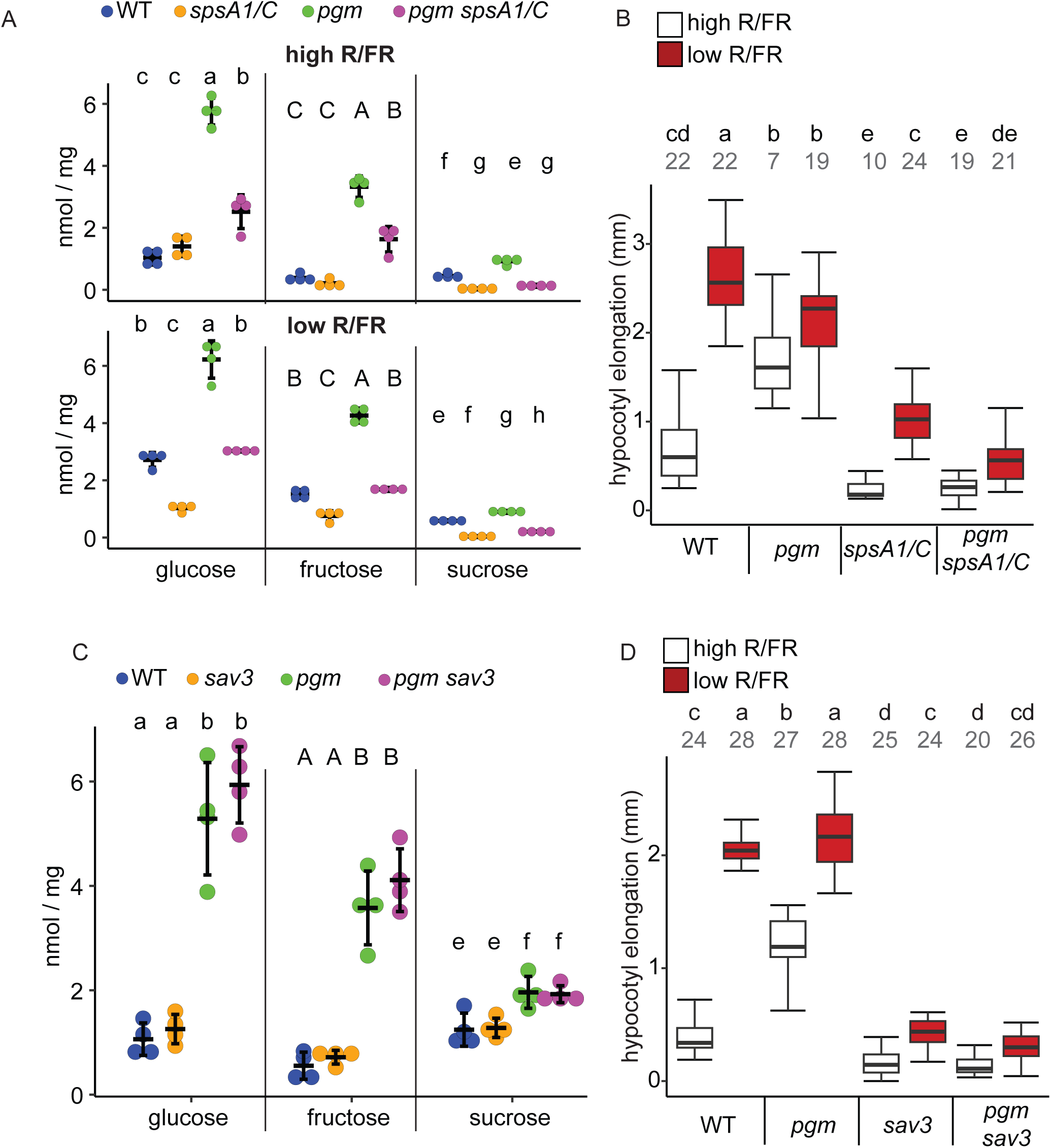
Hypocotyl elongation in the *pgm* mutant requires sucrose and auxin synthesis. A,C) Fructose, glucose and sucrose concentrations from whole-seedling extracts from WT, *pgm*, *spsA1/C* and *pgm spsA1 spsC* (A, 7d-old) or WT, *pgm, sav3* and *pgm sav3* (C, 8d-old). In (A), measurements were done in high (upper plot) and low R/FR (lower plot). Error bars: standard deviation. n=4. B,C) Hypocotyl elongation between 5d and 8d in high or low R/FR in WT, *pgm, spsA1/C* and *pgm spsA1 spsC* (B) or WT, *pgm, sav3* and *pgm sav3*. Grey numbers (B,D) indicate number of hypocotyls measured. Different lower-case letters (A-D) indicate significant differences (ANOVA followed by Tukey-HSD test, P<0.05).

**Figure 3:**
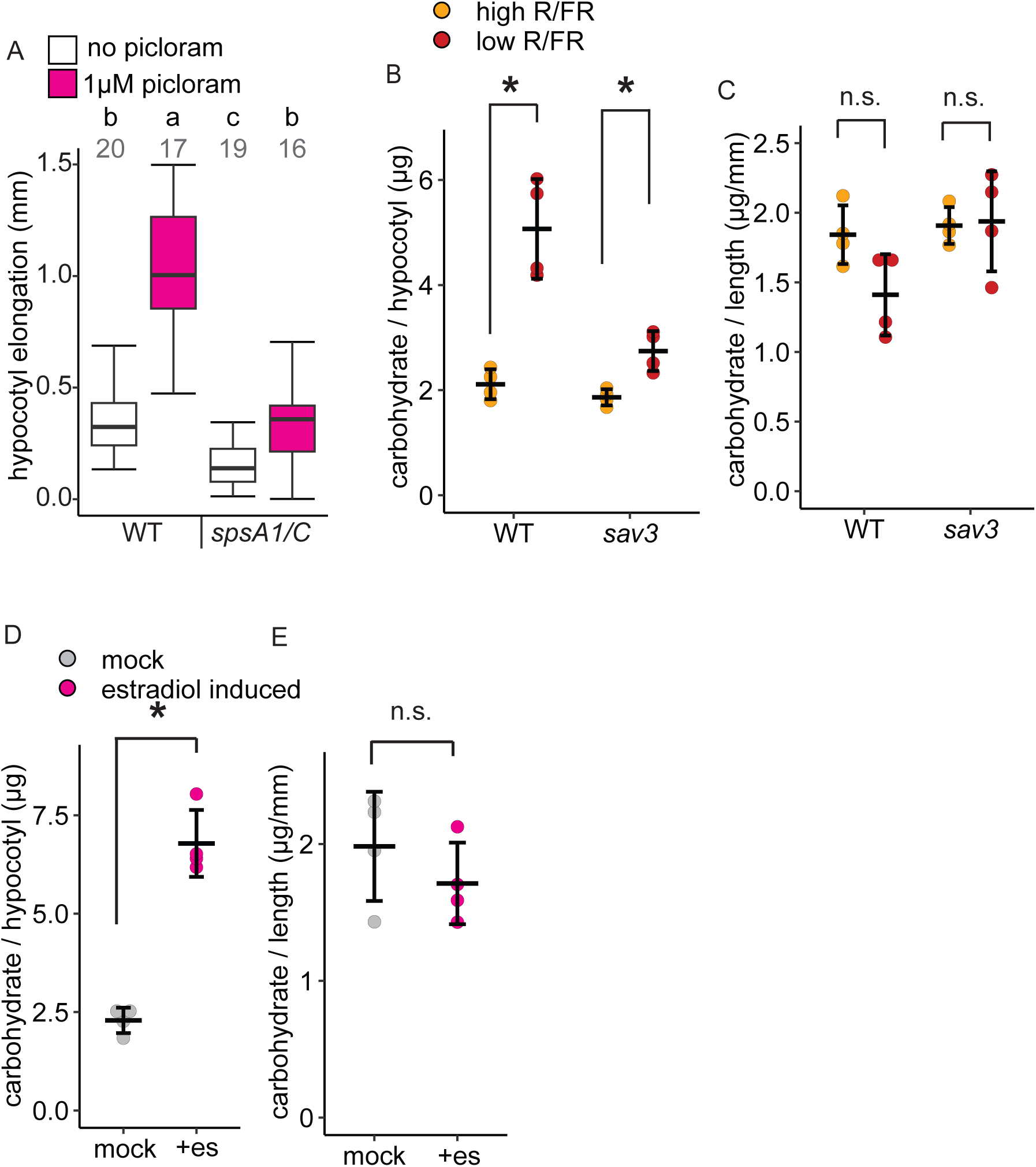
Auxin is necessary and sufficient for biomass partitioning to the hypocotyl and requires sucrose for causing elongation. A) Hypocotyl elongation of WT and *spsA1/C* seedings in response to picloram treatment. Different letters indicate statistically significant difference by Tukey HSD-test, performed after significant (P<0.05) outcome of ANOVA. Grey numbers indicate number of seedlings analysed. B-E) Carbohydrate content of 8d-old Arabidopsis hypocotyls measured with the anthrone assay and expressed as μg glucose-standard equivalent per hypocotyl (B, D) or per mm hypocotyl length (C,E), n=4, where each of the 4 replicates is the average of 10 to 20 hypocotyls. B,C) WT and *sav3-2* hypocotyls with the last 3 days in either high or low R/FR. D,E) Line XVE::FRO6::YUC3 with and without induction with estradiol for the last 3 days before sampling, es = estradiol.* statistically significant by Student’s T-test (B-E). n.s. = not significant by Student’s T-test (C, E).

**Figure 4:**
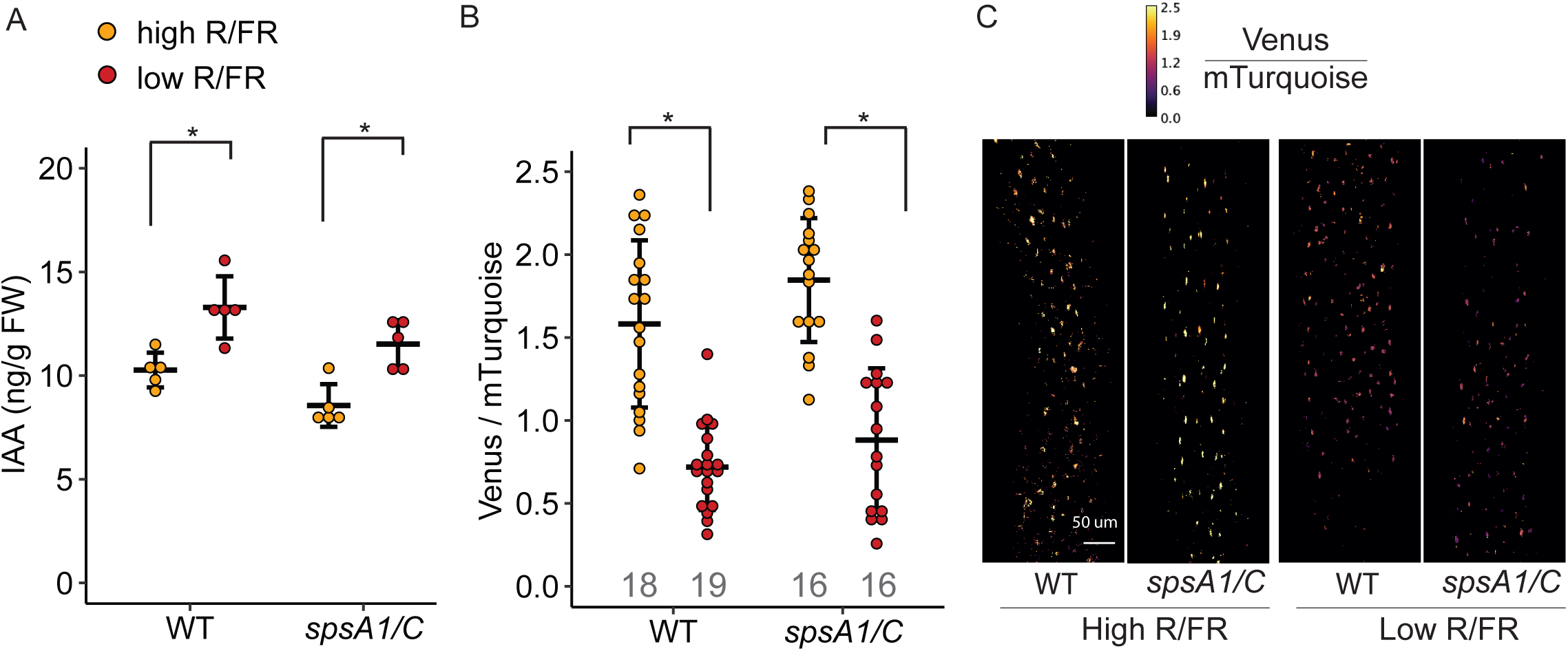
Auxin synthesis induction is not impaired in *spsA1/C* in low R/FR. (A) auxin quantification by LC/MS in WT and *spsA1/C* extracts of 6d-old seedlings after 2h in high or low R/FR. n=5 (B) Quantification of auxin signaling using the fluorescent ratiometric auxin signaling reporter qDIIV (pPDF1::DII-n7-Venus-2A-mTurquoise-sv40) in WT and *spsA1/C* hypocotyls of 5-day old seedlings after 2h of high or low R/FR exposure. Number of replicates are indicated by grey numbers. (C) Representative microscopy images of qDIIV in WT and *spsA1/C* showing the ratio of intensities of Venus to mTurquoise. Scale bar and colour scale applies to all four images. *P<0.05 (t-test). A, B: mean and error plots, error bars: standard deviation

**Table 1:** Characteristics of all light sources (measured by PG200n (UPRTek), intensities in µmol / (m^2^*s). Estimated phytochrome activation state was calculated using transition spectra (Mancinelli, 1994) (P_fr_/P_tot_).

|  | high R/FR<br>(Percival) | low R/FR<br>(Percival) | low blue (Fig.<br>6B), Percival | high R/FR<br>(Polyklima) | low R/FR<br>(Polyklima) |
| --- | --- | --- | --- | --- | --- |
| PAR, in $\mu\text{mol} / (\text{m}^2 \cdot \text{s})$ | 50.5 | 50.4 | 33 | 51.55 | 51.31 |
| red (640-700nm), $\mu\text{mol} / (\text{m}^2 \cdot \text{s})$ | 2.6 | 2.8 | | 8.465 | 8.814 |
| far red (700-760nm), $\mu\text{mol} / (\text{m}^2 \cdot \text{s})$ | 2.0 | 16.4 | | 2.64 | 15.92 |
| R/FR | 1.3 | 0.17 | 1.3 | 3.21 | 0.55 |
| calculated $P_{fr}/P_{tot}$ | 0.79 | 0.60 | | 0.77 | 0.61 |

For all experiments except leaf elevation angle measurements in Arabidopsis, seeds were surface sterilized and subsequently plated on half MS plates with either 0.8% agar (horizonal growth for auxin quantification (Fig. 4A), gene expression measurement (Fig. 5, Fig. S4, S5), sugar treatments (Fig. 1E) and esculin experiments (Fig. S2F-G)) or 1.6% agar and nylon meshes (hypocotyl elongation assays, qDIIV (Fig. 4B-C)). After 4 or 5 days of stratification, plates were moved to an incubator with white light (high R/FR) and 21°C, at Zeitgeber time (ZT) 2.

**Figure 5:**
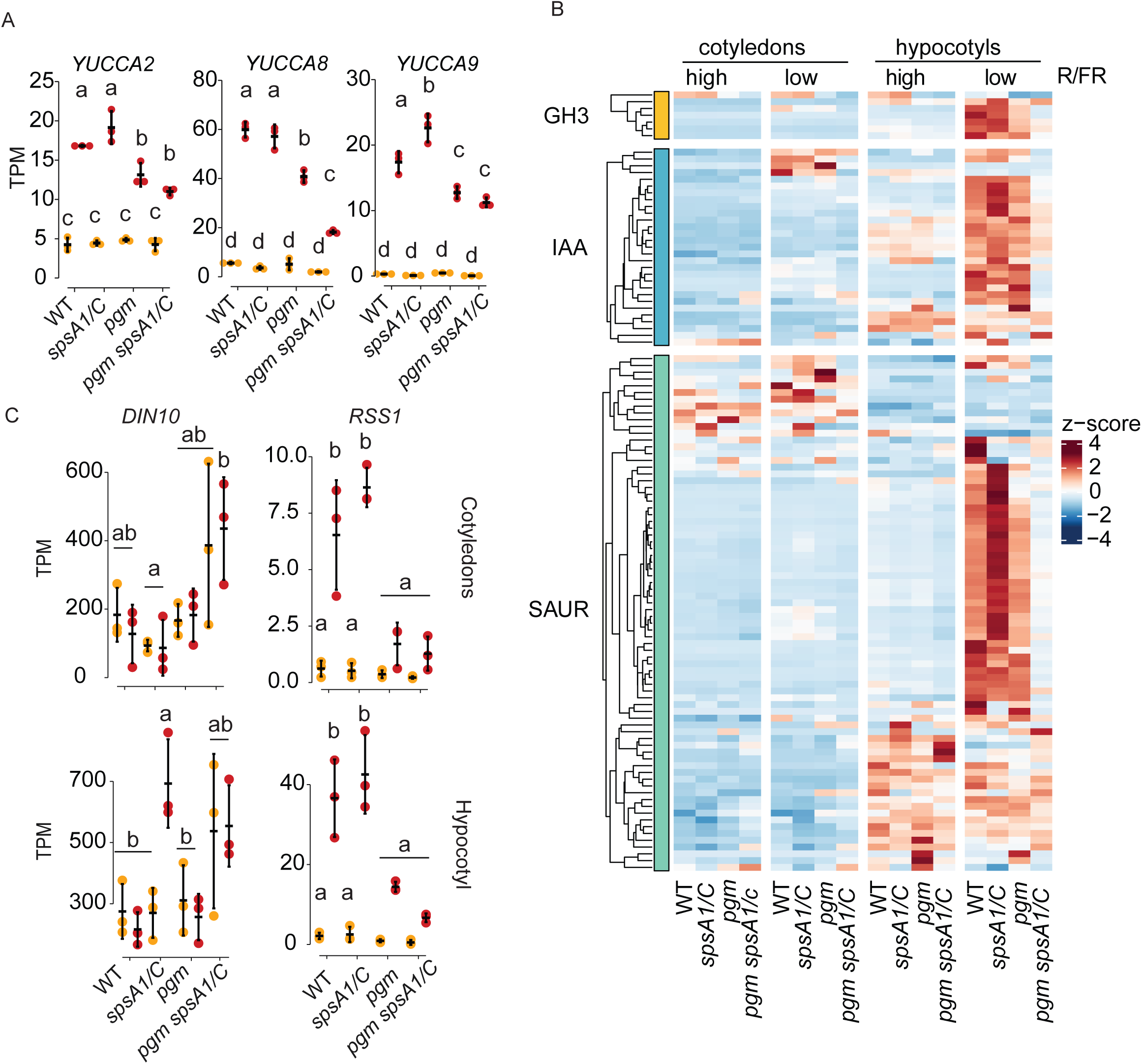
Transcriptomics analysis of cotyledons and hypocotyls in high and low R/FR of WT, *spsA1/C, pgm* and *pgm spsA1/C*. A) Expression of the three *YUCCA* genes which had the strongest low R/FR response in WT cotyledons. B) Heatmap of z-scores of auxin signaling genes of the GH3, IAA and SAUR families. C) Expression of the starvation marker gene *DIN10* and the gene *RSS1*. A,C: mean and error plots, error bars indicating standard deviation. Different letters indicate statistically significant difference (Tukey HSD test, P<0.05). n=3.

For leaf elevation angle measurements, plants were stratified without prior surface sterilization and sown directly on soil (Michaud *et al*., 2017).

### Hypocotyl elongation assays

For hypocotyl elongation experiments, plates were either kept in control conditions (high R/FR without pharmacological treatments) or moved to treatment conditions (different light input, temperature or pharmacological reagent) after 5 days. For experiments with estradiol induction or pharmacological treatments, Arabidopsis seedlings on meshes were transferred to new plates with the treatment substance or only the solvent background (‘mock’), at ZT2 on the 6^th^ day (picloram at 1µM, estradiol at 5µM to induce FRO6::XVE:YUC3). For sucrose treatment (Fig. 1E), 0.75µl of either 90mM sucrose or 90mM sorbitol were applied to the more level cotyledon of each seedling on day 6, 7, and 8.

*Brassica rapa* seedlings were grown on vertical plates without meshes in otherwise the same conditions as Arabidopsis seedlings. For pharmacological treatments with picloram or PPBO, seedlings were transferred one by one to mock or vertical plates on the 5^th^ day of growth (PPBO, at 10, 30 or 100µM (Fig. S3A)) at ZT8, i.e. the evening before the start of changing light conditions, or on the 6^th^ day at ZT2 (picloram, 5µM, Fig. S3B).

Arabidopsis hypocotyl lengths were measured from photographs on plates after 5d and after 8d of growth. Length of each hypocotyl was measured from photos with ImageJ (Schneider *et al*., 2012) and the difference in length between the two time points was calculated for each seedling. Genotypes were blinded during experimentation and hypocotyl measurements.

Brassica hypocotyls were photographed after 8d of growth, from photographs of cut hypocotyls.

### Hypocotyl DW measurement

For hypocotyl biomass measurements, hypocotyls were photographed, dissected and fresh weight was determined immediately. Brassica hypocotyls were dried in aluminum foil in groups of 4 hypocotyls and weighed on a standard lab microbalance after 4 days of drying at 80°C. Hypocotyl dry mass was calculated by dividing the DW by the number of hypocotyls. Arabidopsis hypocotyls were dried in groups of 50 to 100 on small Teflon plates to avoid sticking to the surface so that hypocotyls could slide into specialized weight boats for weighing on an ultra-microbalance (XPR2U, Mettler-Toledo).

### Carbohydrate content by anthrone assay

For each replicate, 10 to 20 hypocotyls of 8-day old seedlings were dissected, homogenized with a plastic pestle in 50µl distilled water, and hydrolyzed by adding 100µl concentrated sulfuric acid and 2h incubation at room temperature. 30µl of the hydrolysate were added to 400µl anthrone reagent (0.2% w/v anthrone in concentrated sulfuric acid) and heated at 80°C for 30min. Glucose standards were incubated with anthrone in the same manner. Absorbance at 620nm was measured on a plate reader and the standard curve of glucose standards was used to calculate carbohydrate content, expressed in µg glucose standard equivalent / hypocotyl. Carbohydrate content / hypocotyl length and DW / fresh weight was determined as described for the DW measurements.

### Quantification of sugar concentrations

The concentrations of glucose, fructose and sucrose in plant extracts were measured in extracts from 7 or 8d-old seedlings using an enzyme-based assay (Cross *et al*., 2006). 30 to 60 whole seedlings were quickly sampled, weighed, flash-frozen in liquid nitrogen and milled by a TissueLyzer (Qiagen). An ethanolic extract was prepared by extracting twice with 80% ethanol / 20% 10mM HEPES/KOH pH7.0 (first with 250µl, second extraction with 150µl) for 20min at 80°C, followed by a third extraction with 250µl 50% ethanol / 50% 10mM HEPES/KOH pH7.0. To measure concentrations of glucose, fructose and sucrose, 30µl of the pooled extracts were added on a 96 well plate with 160µl assay mix (100mM HEPES/KOH pH7.0, 0.86M ATP (Roche), 0.3M NADP (Roche), 3.4U/ml G6PDH (grade II from yeast, Roche). Absorbance at 340nm was measured with sequential addition of 2µl of a 900U/ml hexokinase (Roche) solution, 2µl of a 600U/ml phosphoglucose isomerase solution (from yeast, Roche), and 2µl of a saturated invertase (Sigma-Aldrich) solution. Enzymes were resuspended or dissolved in in 10mM HEPES/KOH pH 7.0. Each enzyme was added after stabilization of absorbance. Glucose concentration in the solution was determined by the absorbance difference caused by addition of hexokinase, fructose by the addition of phosphoglucose isomerase, sucrose by the addition of invertase. Standards on the same plate were used to make a standard curve for quantification.

### Esculin experiments

Esculin was applied to 6-day old seedlings grown high R/FR followed by 2h of high or low R/FR. A 1µl drop of 10mM Esculin in water (65°C to dissolve) was placed onto a cotyledon with a small cut, which did not rupture the mid vein. Seedlings were returned to the same conditions as before. After 25min, cotyledons were blotted to remove external esculin, and seedlings were pulled out of the agar, washed with distilled water and mounted on microscopy slides. Esculin was imaged on a Zeiss LSM 710 confocal miscrope, with excitation at 405nm and emission wavelengths 462nm to 517nm.

Experiments were set up in several batches such that only up to 8 seedlings were analysed at the same time and on one slide, because we observed that the esculin signal decreases slightly over time after mounting. Genotypes and conditions were blinded to the microscopist.

The esculin signal was quantified using ImageJ (Schneider *et al*., 2012). For each hypocotyl image, 10 same-size circular ROIs were placed on the vasculature, 10 into the cortex and the average cortex signal was subtracted from the average vasculature signal.

### qDIIV microscopy

5d old seedlings were harvested after 3h of high vs. low R/FR treatment and, in order to preserve the state of the reporter for all samples at the same time point, immediately fixed with 4% paraformaldehyde in 1x phosphate buffered saline using 3 rounds of vacuum application and release, followed by overnight storage at 4°C.

pPDF1:qDIIV was imaged on a Leica Stellaris microscope, with mTurquoise excitation at 440nm, emission at 445nm-480, Venus excitation at 515nm, emission at 524nm-560, 10x dry objective and in photon counting mode. Intensities were quantified with ImageJ (Schneider *et al*., 2012) by selecting 10 nuclei as ROIs in the middle 50% of each hypocotyl in images showing mTurquoise, and measuring intensities of mTurquoise and Venus. The ratio of these intensities was plotted as the qDIIV signal.

### RNA-seq

Arabidopsis seedlings were grown for five days, then either maintained in high R/FR or transferred to low R/FR at ZT2 until ZT5. Each biological replicate consisted of 25–30 seedlings grown on ½ MS plates. Harvested seedlings were flash-frozen in liquid nitrogen, stored overnight at −70°C, then transferred to −20°C overnight in RNA*later*™-ICE solution before being dissected as described previously (Ince *et al*., 2022). Total RNA extracted using the RNeasy Plant Mini Kit (Qiagen) was used to generate libraries with the Illumina Stranded mRNA Prep kit (Illumina). Sequencing was performed on an Aviti sequencer (Element Biosciences).

Raw reads were trimmed using Cutadapt (v2.5; (Martin, 2011)), filtered for ribosomal RNA with fastq_screen (Wingett & Andrews, 2018) (v0.11.1), and further filtered for low-complexity sequences using reaper (v15-065; (Davis *et al*., 2013)). Clean reads were aligned to the *Arabidopsis thaliana* reference genome (TAIR10.39) using STAR (v2.5.3a; (Dobin *et al*., 2013)), and gene counts were obtained with htseq-count (v0.9.1; (Anders *et al*., 2015)). Alignment quality was evaluated using RSeQC (v2.3.7; (Wang *et al*., 2012)). Differential gene expression analysis was performed with DESeq2 (Love *et al*., 2014), applying thresholds of *p*.adj < 0.05 and |log₂FoldChange| > 1. Gene Ontology (GO) enrichment analysis was done using the *compareCluster* function from the *clusterProfiler* package (Yu *et al*., 2012).

### Auxin quantification by LC-MS

Growth conditions for quantification of free indole-3-acetic acid (IAA) was as in hypocotyl elongation assays in the corresponding light condition but low R/FR treatment (ZT2) and harvesting (ZT5) was done after 6 full days of growth to ensure enough plant material was available. 5 replicates were analyzed for each genotype and condition and each replicate consisted of 300 entire seedlings, amounting to about 200mg. Samples were immediately weighed and frozen in liquid nitrogen. Free IAA was measured by the analysis service lead by Dr Gaetan Glauser at the University of Neuchatel by LC-MS (Glauser *et al*., 2014).

### Leaf elevation angle measurements

Experimental conditions and procedures for leaf elevation measurement of WT and *spsA1/C* plants were as described previously (Michaud *et al*., 2017).

#### Quantification and statistical analysis

We used Student’s T-test when comparing two samples, and ANOVA followed by Tukey’s HSD test when comparing three or more samples. We regarded differences as statistically significant where P-values were below 0.05. We used R (R Core Team, 2021) for all data analyses apart from the leaf angle experiment (Fig. 6C-D), where we used matlab. In mean-and-error plots, error bars represent the standard deviation. In boxplots, the middle bar indicates the median, the box surrounds the data in the second and third quartiles, whiskers show the data range.

**Figure 6:**
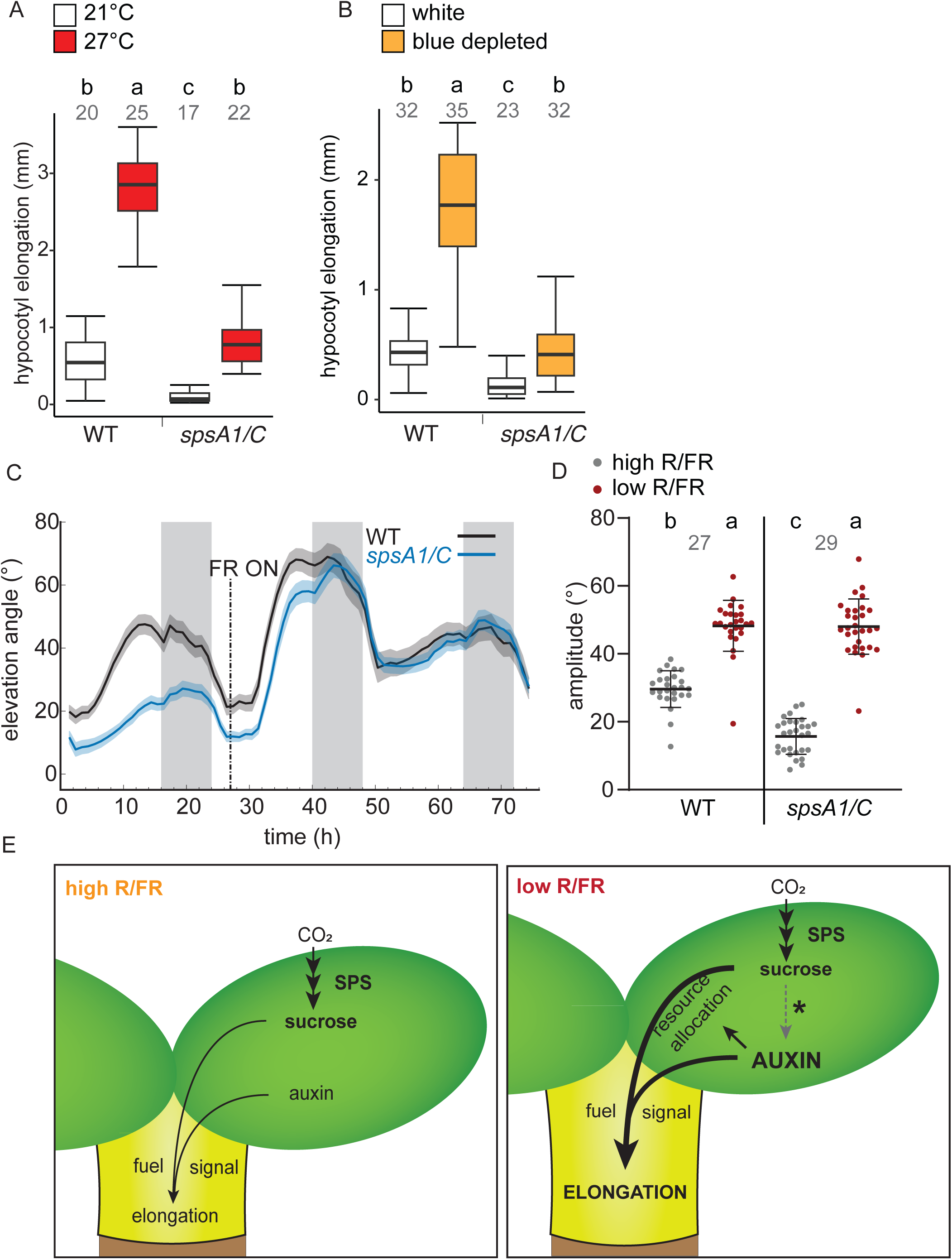
Requirement of SPS for other auxin mediated processes which differ in their growth requirements. A, Hypocotyl elongation after 5d at 21°C followed by 3d at 21°C (white) or 27°C (red). B) Same as (A) but comparing white light vs. blue depleted light (yellow) during the last 3 days of growth. (C) Time course of leaf elevation angles of 2-week old rosettes of WT and *spsA1/C* in high R/FR for 1 day followed by 2 days in low R/FR (from ZT3). D) Quantification of daily leaf elevation amplitudes from time courses shown in (C). E) Working model on the roles and interactions of auxin and sucrose for hypocotyl elongation. Auxin causes resource allocation to the hypocotyl, and both sucrose and auxin cause elongation but require each other. *Sucrose can under some circumstances enhance auxin or auxin signaling, which is supported by Sairanen *et al*. 2012, Stewart et al 2011, Lilley et al., 2012, Singh et al., 2017, de Wit et al 2018 (*pgm pif7*) and Fig. 2D (*pgm sav3*), but not reflected by transcriptional patterns (Fig. 5, Fig S4, S5). Grey numbers in (A), (B) and (D) indicate number of replicates.

## Results

### Sucrose synthesis capacity determines hypocotyl elongation in the neighbor detection response

It is unknown how phytochrome signaling intersects with sugar allocation during the neighbor proximity response. Many processes are involved in sugar allocation and could therefore be subject to regulation, for example sugar transport (de Wit *et al*., 2018). We reasoned that even an earlier process could control hypocotyl elongation via regulating the availability of building blocks and fuel: the sucrose biosynthetic pathways in the source organs, the cotyledons in young seedlings.

We therefore tested whether mutants of enzymes which are critical for sucrose biosynthesis, have an altered hypocotyl elongation response to neighbor proximity simulated by addition of far-red light (Fig. S1A,B). Two biosynthetic steps have been proposed as the key regulatory points of the pathway (Fig. 1A): Firstly, dephosphorylation of fructose-1,6-bisphosphate to form fructose-6-phosphate, catalyzed by cytosolic fructose-bisphosphatase (CYFBP), directing carbon towards sucrose or cell wall synthesis as opposed to starch synthesis. Secondly, production of sucrose-6-phosphate (S-6-P) from UDP-glucose and fructose-6-phosphate, catalyzed by sucrose-phosphate synthase (SPS) (Strand *et al*., 2000).

*cyfbp* mutant plants are viable without major growth impairment due to a plastidial fructose-1,6-bisphosphatase isoform (Rojas-González *et al*., 2015). *cyfbp* mutant seedlings had only slightly shorter hypocotyls than the WT in low R/FR (Fig. 1B), suggesting that other F-6-P sources such as the oxidative pentose phosphate pathway or a fructose-1,6-bisphosphatase located in the chloroplasts (Rojas-González *et al*., 2015) can provide fructose-6-phosphate for sucrose formation in cotyledons. CYFBP is therefore unlikely to be a major point of regulation in response to low R/FR. We also tested a mutant of the enzyme 6-phosphofructo-2-kinase/fructose-2,6-bisphosphatase (F2KP). F2KP produces and breaks down the low-abundance metabolite fructose-2,6-bisphosphate, which inhibits CYFBP (Nielsen *et al*., 2004). Loss of F2KP causes higher sucrose and lower starch levels because of lower CYFBP inhibition (McCormick & Kruger, 2015). In agreement with its function, the *f2kp* mutant had significantly longer hypocotyls, but, like *cyfbp*, did not have a quantitatively large effect (Fig. 1C).

Arabidopsis contains four SPS isoforms, *SPSA1, SPSA2, SPSB* and *SPSC*. We tested the hypocotyl elongation of all mutant combinations which are able to complete the life cycle (Figure 1D, Fig. S1F) (Bahaji *et al*., 2015). The only mutant with a substantial reduction in hypocotyl elongation was the *spsA1 spsC* double mutant (‘*spsA1/C*’) (Fig. 1D). Application of 90mM sucrose to the cotyledons of *spsA1/C* during the three days in high or low R/FR fully rescued its hypocotyl length (Fig. 1E). We therefore conclude that a high capacity for sucrose synthesis, via SPSA1 and SPSC, is crucial for neighbor-proximity induced hypocotyl elongation.

To verify that the *spsA1/C* mutant has reduced sucrose concentrations in our experimental conditions, we measured sucrose, glucose and fructose in high and low R/FR (Fig. 2A). Under high R/FR, sucrose was significantly less abundant in s*psA1/C* than in WT, under low R/FR, all three sugars had lower concentrations in *spsA1/C* than in WT (Fig. 2A).

### Elevated endogenous sucrose or hexoses can enhance hypocotyl elongation, requiring auxin

We next asked how hypocotyl growth responds to excess sucrose. Fig. 1E indicates that exogenous sucrose application enhances hypocotyl length of WT seedlings only in low R/FR, but not high R/FR. This observation contrasts with the phenotype of the sugar accumulating *pgm1-1* (‘*pgm*’) mutant, which is substantially longer than the WT in high R/FR (de Wit *et al*., 2018). *pgm* is a loss-of-function mutant of the starch biosynthetic enzyme phosphoglucomutase. Instead of starch, *pgm* plants redirect carbon to high concentrations of sugars (Gibon *et al*., 2004). To confirm that the long hypocotyls in *pgm* are indeed due to its high sugar abundance, we crossed the *pgm* mutant with the *spsA1/C* mutant. The concentration of sucrose of the resulting triple mutant *pgm spsA1/C* is low, resembling the *spsA1/C* mutant, while glucose and fructose concentrations are similar to WT (low R/FR) or slightly higher (high R/FR) (Fig. 2A). The hypocotyls of *pgm spsA1/C* were also as short as or even shorter than the *spsA1/C* mutant (Fig. 2B), indicating that high concentrations of sucrose and / or hexoses are responsible for the *pgm* phenotype. Therefore, high endogenous sugar concentrations, but not exogenous sucrose treatment, can enhance hypocotyl elongation in the absence of a low R/FR cue.

According to a previous study (de Wit *et al*., 2018), the *pif7 pgm* double mutant has the same hypocotyl length in low R/FR as *pif7* in spite of having elevated sugar concentrations like *pgm*. This observation indicates that elevated endogenous sugar levels do not cause elongation if plants are unable to induce the PIF7-dependent neighbor detection signaling pathway. To further narrow down the signaling required for translating high sugar into hypocotyl elongation in *pgm*, we wanted to test whether auxin biosynthesis is also required for its longer hypocotyls in high R/FR. To do so, we generated the *pgm sav3* double mutant. *SAV3* encodes the enzyme TAA1 in the canonical auxin biosynthetic pathway (Tao *et al*., 2008). TAA1 synthesizes the substrate of YUCCA (Tao *et al*., 2008). Even though TAA1 itself is not up-regulated by low R/FR (Tao *et al*., 2008), its loss-of function prevents the increase in auxin concentration in response to low R/FR as the substrate of YUCCAs is unavailable. *pgm sav3* has elevated glucose, fructose and sucrose levels like *pgm* (Fig. 2C) but is unable to elongate in response to low R/FR (Fig. 2 D), behaving like *pif7 pgm* (de Wit *et al*., 2018). Therefore, we conclude that biosynthesis of at least basal auxin levels is required for translating elevated sucrose or hexoses into hypocotyl elongation growth.

### Auxin regulates resource allocation and requires SPS enzymes

Given our results that auxin is required for elongation caused by high sugar levels, we hypothesized that auxin plays a role in resource allocation. Therefore, we tested whether a high sucrose biosynthetic capacity is necessary also for hypocotyl elongation in response to treatment with the auxin analog picloram. Like indole-3-acedic acid (IAA), picloram induces hypocotyl elongation, and activates auxin signaling by a subset of the known IAA receptors (Savaldi-Goldstein *et al*., 2008; Torii *et al*., 2018). Indeed, hypocotyl elongation in response to picloram treatment was strongly impaired in *spsA1/C* (Fig. 3A).

Next, we wanted to test whether auxin is responsible for the resource allocation changes observed in response to low R/FR. So far, evidence for enhanced biomass allocation to the hypocotyl in low R/FR has been given for *Brassica rapa* (de Wit *et al*., 2018) but not Arabidopsis, where hypocotyls are much smaller. We solved this methodological challenge by drying a large number of Arabidopsis hypocotyls, sampled at the end of an elongation assay using a high sensitivity microbalance. The approximately 2-fold hypocotyl length increase in low R/FR (Fig. S2A) was accompanied by a significant increase in DW (Fig. S2B). Therefore, the biomass increase observed in *Brassica rapa* in response to low R/FR (de Wit *et al*., 2018) also applies to Arabidopsis and is largely proportional to length (Fig. S2C).

In order to reduce the number of dissected hypocotyls needed for each experiment, we tested whether the DW measurements correlate well with carbohydrate content measurements using the anthrone assay (Turula *et al*., 2010), which only requires between 10 and 20 hypocotyls per sample. Our rationale is that carbohydrates – found in cell wall, starch and sugars – constitute by far the largest part of the plant’s dry biomass. The results from the anthrone fully agreed with the DW measurements (Fig. S2A-C), confirming that the anthrone assay is a good proxy for hypocotyl DW. We used the anthrone assay to compare the WT, *spsA1/C* and *pgm* mutants and found that *pgm* hypocotyls contain more, *spsA1/C* less carbohydrate than WT (Fig. S2D). Notably, the *spsA1/C* hypocotyls had a lower carbohydrate content per length (Fig. S2E), indicating that this mutant achieves the small amount of growth at a slightly lower carbon density.

We then tested whether auxin induces resource allocation changes in Arabidopsis. First, we tested if auxin is necessary for enhancing hypocotyl biomass in low R/FR. Indeed, the *sav3-2* mutant, which does not have an increase auxin content in low R/FR (Tao *et al*., 2008), had a strongly reduced increase in hypocotyl carbohydrate content in low R/FR compared with the WT (Fig. 3B), proportionally to hypocotyl length (Fig. 3C). To test whether induction of auxin synthesis is sufficient for carbon allocation to the hypocotyl, we made use of the line XVE::FRO6::YUC3 which allows inducible expression of the auxin bottleneck biosynthetic gene *YUC3* in the cotyledons (Chen *et al*., 2014). Upon induction with estradiol, hypocotyl dry biomass increased approx. 3-fold (Fig. 3D), proportionally to hypocotyl length (Fig. 3E).

Because carbon is transported between organs as sucrose in the phloem, we predicted that increased carbon allocation to the hypocotyl is accompanied by increased sucrose transport from cotyledons to the hypocotyl. To test this early sign of changes of resource allocation, we tested the transport of the phloem-mobile dye esculin to the hypocotyl after 3 hours of high vs. low R/FR treatment. Esculin is loaded into the phloem by the SUC2 transporter, and subsequently travels from source to sink following the phloem mass flow (Knox *et al*., 2018). To test whether esculin enters the phloem via SUC2 in cotyledons, we loaded a drop of esculin onto cotyledons of WT and *suc2-4* mutant plants which express a functionally impaired SUC2 transporter (Srivastava *et al*., 2008). While esculin stained the vasculature in the WT, it only stained the surface of cotyledons in *suc2-4* (Fig. S2F). We therefore proceeded to quantifying the esculin signal in the vasculature of hypocotyls, after 3h of high vs. low R/FR of WT. We also included *sweet11/12* mutants, which have a reduced capacity for phloem transport and shorter hypocotyls in low R/FR (de Wit *et al*., 2018). While the esculin signal was significantly higher in low than high R/FR in the WT, it was generally lower in *sweet11/12* and the trend for an increase in transport was not significant (Fig. S2G). We conclude that the change in biomass allocation observed in response to low R/FR is noticeable as an early change in sugar phloem transport.

Next, to find out if our observations are more broadly applicable, we tested if auxin also controls resource allocation in *Brassica rapa* (Figure S3). Treatment with the auxin biosynthesis inhibitor p-phenoxyphenyl boronic acid (PPBO) (Kakei *et al*., 2015) inhibited the increase in hypocotyl dry weight (DW) in low R/FR in a dose-dependent manner (Fig. S3A), demonstrating that auxin synthesis is necessary for resource allocation to the hypocotyl. Moreover, picloram treatment significantly enhanced hypocotyl DW in high R/FR (Fig. S3B), showing that an auxin analog is sufficient for enhancing biomass allocation.

Collectively, these experiments indicate that increased auxin production in cotyledons/young leaves is necessary and sufficient to promote allocation of carbon resources to the hypocotyl in *Brassica rapa* and Arabidopsis.

### Auxin concentration and signaling is largely unaffected by reduced sucrose biosynthetic capacity

Given that external glucose or sucrose application can induce auxin biosynthesis in other experimental conditions (Sairanen *et al*., 2012), we asked whether auxin concentrations are altered in *spsA1/C*. Therefore, we measured extracted IAA concentrations in whole seedlings in WT and *spsA1/C* 3h after moving seedlings into different light conditions. We found that IAA concentrations in *spsA1/C* responded similarly to the WT (Fig. 4A).

Since the IAA measurements were done from whole-seedling extracts, where cotyledon tissue dominates, we decided to use a genetically encoded auxin signaling input sensor to image auxin in hypocotyls. We used the reporter pPDF1::DII-n7-Venus-2A-mTurquoise (pPDF1:qDIIV) (Galvan-Ampudia *et al*., 2018; Boccaccini *et al*., 2020), where Venus but not mTurquoise is subject to degradation at higher levels of auxin. A lower ratio is therefore indicative of a higher auxin concentration. We measured the Venus / mTurquoise ratio in hypocotyls after 3h of high vs. low R/FR. As in the case of IAA content, the qDIIV ratio responded in the same way in *spsA1/C* as in WT (Fig. 4B-C), indicating that the auxin increase in the hypocotyl was not impaired by the lower levels of sucrose.

To get a bigger picture of the influence of sucrose levels in seedlings on low R/FR-induced regulated signalling, we performed gene expression analyses. We compared four genotypes (WT, *spsA1/C, pgm, pgm spsA1/C*), in two organs (hypocotyls and cotyledons) in both light conditions (high and low R/FR for 3h) (Table S1, Table S2). Principal component analysis (PCA) demonstrated that the light condition is the primary source of transcriptional variation across samples. However, the degree of genotype-dependent separation within each light condition differs between hypocotyls and cotyledons (Figure S4A, B).

The classical auxin-independent shade marker transcription factors *HFR1*, *PIL1* and *ATHB2* were induced by low R/FR in all genotypes (Fig. S4C). Interestingly, all three transcription factors responded even more strongly in *spsA1/C* than the WT, and *HFR1* showed a slightly reduced response in *pgm* than WT. This indicates that endogenous sugar levels have a limited influence on the early transcriptional response to low R/FR. Moreover, GO terms related to auxin and the response to R or FR light were up-regulated in all genotypes in hypocotyls and cotyledons (Fig. S4D, Table S3), indicating that the transcriptional response related to auxin is not globally impaired in *spsA1/C* or *pgm*.

Because induction of auxin synthesis in cotyledons in response to low R/FR is due to transcriptional up-regulation of *YUCCA* genes (Müller-Moulé *et al*., 2016), we were interested in their expression levels. The three *YUCCA* genes which had the strongest transcriptional response in WT cotyledons (*YUC2, 8, 9*) responded normally in *spsA1/C* and were slightly reduced in *pgm* cotyledons, and baseline levels were similar in all genotypes (Fig. 5A). Similarly, the expression of early auxin-response genes of the *GH3*, *IAA* and *SAUR* families in the hypocotyl, did not indicate striking differences between WT, *spsA1/C* and *pgm* (Fig. 5B). However, most low R/FR responsive auxin signaling genes had a substantially reduced response in *pgm spsA1/C* compared to the other genotypes (Fig. 5B, S4C, S5).

Auxin signaling induces hypocotyl elongation by several mechanisms, including regulation of cell wall biosynthetic genes. Behavior of low R/FR responsive cell wall-related genes in the hypocotyl was not identical between mutants, but in the bigger picture, induction was observed in WT, *spsA1/C* and *pgm* (Fig. S5). Here, again the *pgm spsA1/C* mutant had a strongly reduced response. As *pgm spsA1/C* is unable to make starch as well as large quantities of sucrose, we expect its metabolism to be strongly mis-regulated. We therefore tested whether we could detect altered expression of starvation-related genes as one form of metabolic misalignment. Indeed, the starvation marker *DARK INDUCIBLE (DIN) 10 was* induced in *pgm spsA1/C* in high and low R/FR, and in both organs (Fig. 5C).

Notably, *DIN10* was also induced in *spsA1/C* only in hypocotyls and only in low R/FR, suggesting a lack of carbon supply. The specific upregulation of *DIN10* in hypocotyls of low R/FR treated seedlings may be an indication that these seedlings induce the signaling pathways to enhance hypocotyl growth but cannot elongate due to reduced sucrose availability.

The transcription factor REGULATED BY SUGAR AND SHADE1 (RSS1) is a negative regulator of auxin signaling and hypocotyl elongation and has previously been suggested as a link between glucose availability and hypocotyl growth as it is suppressed by elongation-inducing glucose treatment (Singh *et al*., 2017). Indeed, in hypocotyls, *RSS1* expression is slightly reduced in *pgm* and *pgm spsA1/C* (Fig. 5C). However, as the *pgm* mutant does not have an increased expression of auxin synthesis and response genes in our data (Fig. 5A, B), it is unlikely that *RSS1* is critical to connect sugar with auxin homeostasis in our experimental conditions.

Taken together, our transcriptomic analysis confirms our data from direct auxin measurements in *spsA1/C* and from the auxin signaling input reporter in *spsA1/C* hypocotyls (Fig. 4). We thus conclude that reduced hypocotyl elongation in *spsA1/C* is not due to altered auxin biosynthesis and signaling and that despite low sugar content, *spsA1/C* can mount a normal low R/FR-induced auxin response. However, the specific induction of *DIN10* in the hypocotyl of low R/FR treated *spsA1/C* reveals an early sign of carbon starvation in this mutant.

### Sugar limitation impairs other hypocotyl growth responses but not the leaf elevation response

Our results from *spsA1/C* indicate that lack of sucrose prevents hypocotyl elongation because of the lack of carbon resources, rather than because of inhibition of the phytochrome and auxin signaling pathway. To challenge this conclusion, we tested the response of *spsA1/C* to other conditions which require rapidly enhanced growth – the hypocotyl elongation response to elevated temperature (Fiorucci *et al*., 2020) (Fig. 6A), and to blue light depletion (Keller *et al*., 2011; Pedmale *et al*., 2016; Ince *et al*., 2022) (Fig. 6B). In both cases, *spsA1/C* had a reduced elongation response, similar to its impairment in low R/FR. By contrast, *spsA1/C* had a strong leaf elevation response to low R/FR (Fig. 6C), which may reflect that only small amounts of growth, differing for ab-and adaxial side of the petiole, are needed for a hyponastic response.

## Discussion

Our study shows that SPSA1 and SPSC are essential for hypocotyl elongation, most likely because they supply the sucrose required to sustain growth. This requirement is evident across multiple elongation-inducing cues, including neighbour detection (Fig. 1D), picloram treatment (Fig. 3A), elevated temperature (Fig. 6A), and blue-light depletion (Fig. 6B). Importantly, reducing sucrose biosynthetic capacity did not impair the low-R/FR-induced increase in auxin accumulation or the associated transcriptional reprogramming (Fig. 4, Fig. 5; Fig. S4, S5). This is consistent with the short-hypocotyl phenotype of *spsA1/C* being due to insufficient carbon allocation to the hypocotyl rather than altered auxin production or signalling.

Despite this strong requirement for endogenous sucrose production, exogenous sucrose does not induce hypocotyl elongation in WT seedlings in the absence of a neighbour-detection signal (Fig. 1E). In contrast, the *pgm* mutant, which accumulates high endogenous sucrose and hexose levels, does elongate under high R/FR ((de Wit *et al*., 2018), Fig. 2B,D). Thus, the conceptual role of sucrose differs from that of auxin: auxin is both necessary and sufficient for hypocotyl elongation, including when supplied exogenously (Tao et al., 2008), whereas sucrose is necessary but not sufficient. Here we show that auxin is necessary and sufficient to direct carbon resources towards the hypocotyl (Fig. 3B–E; Fig. S3). Because low R/FR triggers a localized increase in auxin production and carbon export only from the treated leaf (Casal *et al*., 1995; Michaud *et al*., 2017; Pantazopoulou *et al*., 2017), we propose that elevated auxin in the leaf or cotyledon exposed to low R/FR may initiate carbon mobilization (Fig. 6E). Together, these results lead us to conclude that hypocotyl elongation under low R/FR requires both auxin and sucrose transport to the hypocotyl, with auxin in the photosynthetic tissues triggering allocation of enhanced carbon resources towards the growing hypocotyl (Fig. 3; Fig. 6E; de Wit et al., 2018).

Effects of sugars on signaling pathways can indicate that sugars function as signaling molecules in addition to their role as carbon resources (Ljung *et al*., 2015). Therefore, the question arises whether sucrose or hexoses have such a dual function in the neighbor detection response. Our data demonstrate that sucrose is required for hypocotyl elongation (Fig. 1D-E). Combined with the enhanced expression of the carbon starvation marker gene *DIN10* (Peixoto *et al*., 2021) exclusively in *spsA1/C* hypocotyls and low R/FR (Fig. 5C), it is highly plausible that sucrose acts as a carbon resource to fuel hypocotyl elongation. Two other starvation markers, *DIN6* and *BCAT2*, had increased expression in *pgm spsA1/C* but did not show the low R/FR and hypocotyl specific increase in *spsA1/C* (Table S1, Table S2). Interestingly, *DIN6* and *BCAT2* function in both nitrogen and carbon metabolism, while *DIN10* encodes a carbon metabolic enzyme (a raffinose synthase), possibly indicating a lack of specifically carbon resources in *spsA1/C*.

Whether any of the three sugars sucrose, glucose or fructose also act as a signal regulating hypocotyl growth can currently not be answered because the mechanisms by which elevated sugar in *pgm* operates through *PIF7* and *SAV3* are unknown. Mechanistic links of how sugars affecting phytochrome and auxin signaling have been proposed in other conditions: In response to elevated temperature, sugars can enhance hypocotyl elongation by stabilizing PIF4 (Fan *et al*., 2025); higher CO_2_ concentrations, which may translate into higher carbon availability, enhance the expression of neighbor detection and auxin signaling responsive genes in *Brassica rapa* (Arsovski *et al*., 2018); links may also involve *RSS1*, a transcription factor which suppresses hypocotyl elongation and is inhibited by exogenous sugar (glucose) application (Singh *et al*., 2017). However, the direct mode of sugar action is unknown in all of these cases. Therefore, it remains unclear whether it acts as a direct signal or affects signaling pathways indirectly.

Our study highlights the importance of sucrose biosynthetic capacity (Fig. 1). Hence an interesting question is whether sucrose biosynthetic enzymes could be regulated by low R/FR to control resource allocation. Transcripts of CYFBP, F2KP, SPSA1 and SPSC are not regulated by low R/FR in cotyledons (Kohnen *et al*., 2016). However, F2KP and SPS activity can be regulated by protein phosphorylation (Hartman *et al*., 2023). SPS activity changes in mustard leaves in response to elongation-inducing end of day far red treatment (Yanovsky *et al*., 1995), and in radish leaves in response to far red supplementation (Keiller & Smith, 1989). SPS regulation is therefore an attractive hypothesis also in Arabidopsis. Besides, auxin and neighbor detection have more broadly been shown to affect sugar and starch metabolism (Yang *et al*., 2016; Jia *et al*., 2020; Krahmer *et al*., 2021; Shi *et al*., 2024).

Sucrose transport may also be regulated by auxin to achieve changes in resource allocation, such as via SWEET11, 12 and SUC2 (de Wit *et al*., 2018). Application of auxin to Arabidopsis leaves enhances SUC2 mediated phloem loading by transcriptional up-regulation of *SUC2*, mediated by ARFs (Ren *et al*., 2024), which is however an unlikely scenario for the neighbor detection response as *SUC2* mRNA is not regulated by low R/FR (Table S1 and (Kohnen *et al*., 2016)). Interestingly, SUC2 can also be regulated post-translationally by phosphorylation or ubiquitylation, notably by light intensity (Xu *et al*., 2020). Moreover, auxin can regulate deposition of callose at plasmodesmata and therefore cell-to-cell permeability, to control its own accumulation and transport (Han *et al*., 2014), and possibly also affecting symplasmic sucrose distribution. Taken together, post-translational regulation of sucrose biosynthetic enzymes and / or sucrose transporters by auxin signaling provides a promising hypothesis.

Further study on this topic is critical to understand the mechanisms governing resource allocation regulation – which is not only an unresolved question in fundamental plant biology, but also relevant for crop yield. In poplar, auxin enhances carbon allocation from leaves to the stems and root (Balasubramanian *et al*., 2024), in mustard from leaves to internodes (Casal *et al*., 1995; Yanovsky *et al*., 1995). In tomato auxin is involved in suppressing formation of new flowers to enhance resource allocation to developing fruits (López-Martín *et al*., 2025). Low R/FR treatment increases carbon allocation to tomato fruits (Ji *et al*., 2020), while it reduces the yield of wheat due to prioritization of stem growth (Golan *et al*., 2024) and decreases resource allocation to the root tubers of radish (Keiller & Smith, 1989) and likely sugar beet (Adjesiwor *et al*., 2021; Ballenger *et al*., 2024).

## Acknowledgements

We thank the Cellular Imaging Facility (CIF) at the University of Lausanne and the Center for Advanced Bioimaging (CAB) at the University of Copenhagen for assistance with confocal microscopy, and Prof Staffan Persson (University of Copenhagen) for funding. Furthermore, we are grateful for lab maintenance by Martine Trevisan (University of Lausanne) and Ouda Ogden (University of Copenhagen), and contributions from student helpers Tiphaine-Marie Lainey, Jana Näf, Clémence Nicollerat and Sandro Speranza for seed plating and hypocotyl measurements. This work was supported by the VELUX Stiftung (project number 1455, awarded to CF and JK); a Tremplin grant awarded to JK by the Advisory Committee on Equality, Diversity and Inclusion, and the Equality Office, Université de Lausanne (UNIL); the Faculty of Biology and Medicine, (UNIL) for parental leave support to JK; Swiss National Science Foundation grant 310030_200318 awarded to CF; NNF Research laureate grant no. NNF19OC0056076 awarded by Novo Nordisk Fonden to Prof Staffan Persson (PLEN, University of Copenhagen).

## Competing interests

The authors declare no competing interests.

## Author contributions

SP, BJRH, CF and JK planned and designed the research; SP, BJRH, JH, YCI and JK performed experiments and analyzed data; SP, BJRH, CF and JK wrote the manuscript. SP and BJRH contributed equally.

## Data availability

The RNA-sequencing data generated in this study is available in the Gene Expression Omnibus database (GEO), under accession no. GSE315941 (Paulišić *et al*., 2026).

## Supplemental item titles

### Supplementary figures

**Figure S1: Spectra in growth cabinets, and hypocotyl elongation response to low R/FR of all *sps* mutants.** A-E) Spectra of light sources used. A) High R/FR cool white fluorescent tubes in Percival. B) Same white light as in (A) with FR LEDs (GroLEDs, Percival) C) Same white light source as in (A) but with a double layered yellow filter (LEE filters number 010, medium yellow). D) Cool white fluorescent LEDs in Polyklima cabinet. E) Same light as in (D) with FR LEDs. Light sources (A) and (B) were used for most low R/FR experiments, with the exception of biomass measurements (Fig. 3B-E, S2A-F), sugar measurements (Fig. 2A,C) and qDIIV (Fig. 4B,C), where light sources in (D) and (E) were used. (C) was used for Fig. 6B. F) Hypocotyl elongation response to low R/FR of all *sps* mutant combinations which are able to complete the life cycle, grouped in single mutants, double mutants and triple mutants. All mutants were measured as part of the same experiment, WT data is the same in all three diagrams and therefore shaded in double and triple mutant plots. Statistical analysis of elongation differences was done for all data together. Different letters indicate significant difference, P<0.05, Tukey HSD test). Grey numbers indicate number of hypocotyls quantified per sample.

**Figure S2: Neighbor detection is accompanied by enhanced resource allocation to the hypocotyl in Arabidopis by three independent methods**. A-C): Experimental evidence of hypocotyl biomass increase after 3 days of high vs. low R/FR, using direct DW measurements and carbohydrate content measurement by the anthrone assay. n=4 with 50 to 100 (DW) or 10 to 20 (anthrone assay) hypocotyls per replicate. A) Hypocotyl length of seedlings harvested for both analyses in high (orange) and low (dark red) R/FR. B) DW and carbohydrate content by anthrone assay in high and low R/FR. Carbohydrate content is expressed in μg glucose standard equivalents. C) DW or carbohydrate (anthrone assay) per mm hypocotyl length in high and low R/FR. D) Carbohydrate content of WT, *spsA1/C* and *pgm* hypocotyls by anthrone assay. n=4, with each anthrone assay replicate consisting of 10 to 20 hypocotyls. E) Data from (D) divided by average length of the hypocotyls used. F-G: Measurement of sucrose transport with the phloem mobile dye esculin, which is loaded into the phloem by the SUC2 transporter, therefore the *suc2-4* mutant shows only surface staining, rather than strong fluorescence in the vasculature (F, scale bar: 500μm). G) Quantified fluorescence intensity in the vasculature, with the cortex background subtracted, in high and low R/FR in WT and *sweet11/12*. Grey numbers indicate number of hypocotyls analysed per sample. Statistics in A-C: * P < 0.05 (Student’s T-test). Statistics in D, E, G: different letters indicate significant difference (P<0.05, Tukey’s HSD test). Error bars in all plots: standard deviation.

**Figure S3: Auxin is necessary and sufficient for resource allocation to the hypocotyl in *Brassica rapa*.** A) Hypocotyl dry weight of 8d old *Brassica rapa* seedlings, grown in high R/FR for 5d followed by 3d in low R/FR with and without PPBO treatment. n=5, where each of the five replicates is the combined dry weight of 4 hypocotyls divided by 4. Different letters indicate statistically significant difference by Tukey HSD-test, performed after an ANOVA with significant outcome (P<0.05). B) As in (A), with the last 3d in high R/FR on either mock or picloram supplemented plates. * statistically significant by student’s T-test.

**Figure S4: Additional analysis of RNA-seq dataset.** A-B) PCA of hypocotyl (A) and cotyledon (B) RNA-seq data. C) Expression of the classical neighbor detection-induced shade marker genes *HFR1*, *PIL1* and *ATHB2*. D) Functional enrichment analysis (GO - biological process) of low R/FR responding genes in hypocotyls. pr. = process. m.o. = multicellular organismal. n=3

**Figure S5: Expression of cell wall growth related genes.** Heat map of z-scores of cell-wall related differentially expressed genes (Sénéchal *et al*., 2024).

### Supplementary tables

**Table S1: TPM of genes from the RNA-seq in response to low R/FR.**

**Table S2: Differential analysis of RNA-seq in response to low R/FR.**

**Table S3: Gene ontology enrichment analysis.**

## Notes

https://www.ncbi.nlm.nih.gov/geo/query/acc.cgi?acc=GSE315941

### Competing Interest Statement

The authors have declared no competing interest.

