## Supplementary material for "Internal sugar allocation in response to a shade signal is regulated by concerted action of auxin and sucrose": Figure S1

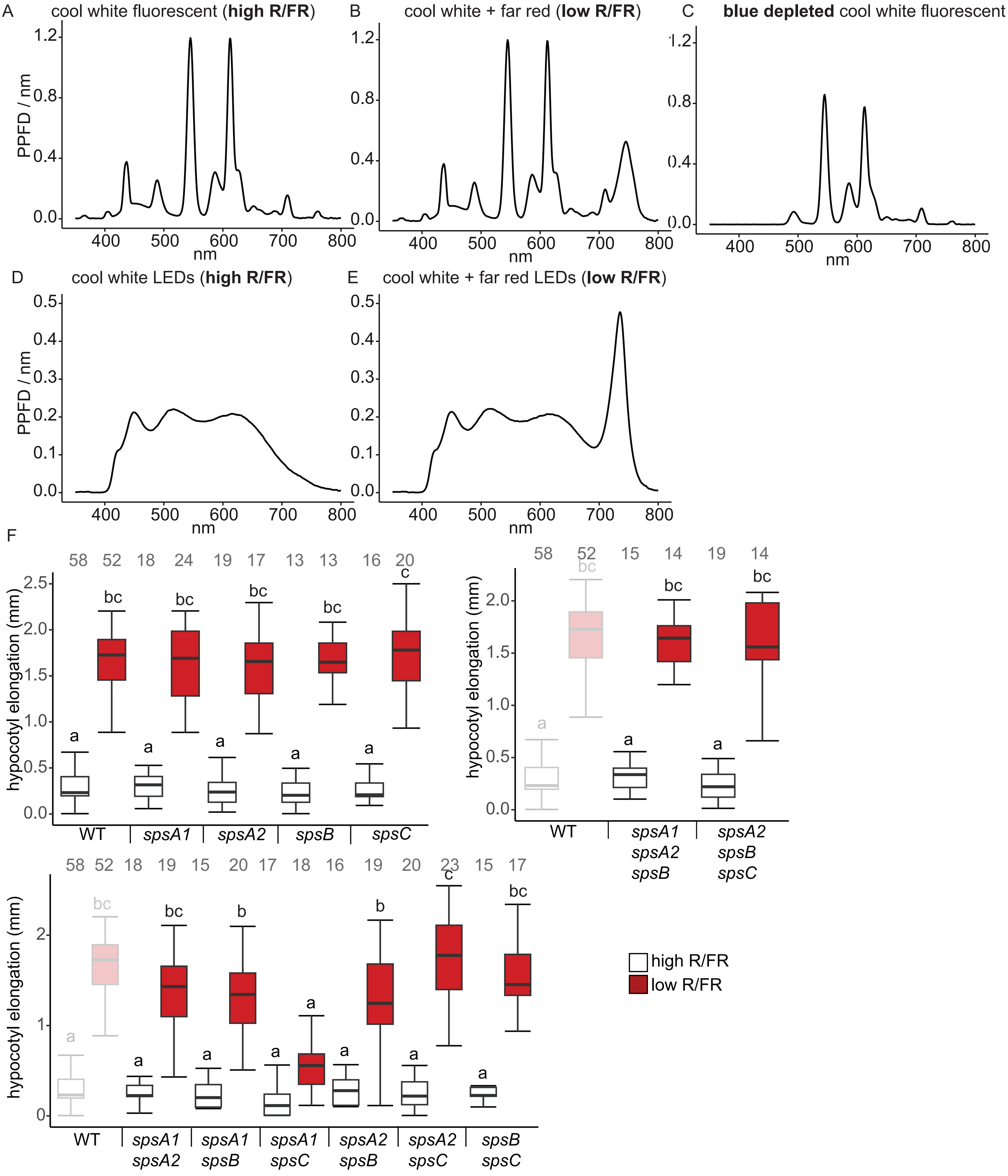


**Figure S1: Spectra in growth cabinets, and hypocotyl elongation response to low R/FR of all *sps* mutants.** A-E) Spectra of light sources used. A) High R/FR cool white fluorescent tubes in Percival. B) Same white light as in (A) with FR LEDs (GroLEDs, Percival) C) Same white light source as in (A) but with a double layered yellow filter (LEE filters number 010, medium yellow). D) Cool white fluorescent LEDs in Polyklima cabinet. E) Same light as in (D) with FR LEDs. Light sources (A) and (B) were used for most low R/FR experiments, with the exception of biomass measurements (Fig. 3B-E, S2A-F), sugar measurements (Fig. 2A,C) and qDIIV (Fig. 4B,C), where light sources in (D) and (E) were used. (C) was used for Fig. 6B. F) Hypocotyl elongation response to low R/FR of all *sps* mutant combinations which are able to complete the life cycle, grouped in single mutants, double mutants and triple mutants. All mutants were measured as part of the same experiment, WT data is the same in all three diagrams and therefore shaded in double and triple mutant plots. Statistical analysis of elongation differences was done for all data together. Different letters indicate significant difference, $P<0.05$, Tukey HSD test). Grey numbers indicate number of hypocotyls quantified per sample.
