## Supplementary material for "Internal sugar allocation in response to a shade signal is regulated by concerted action of auxin and sucrose": Figure S2

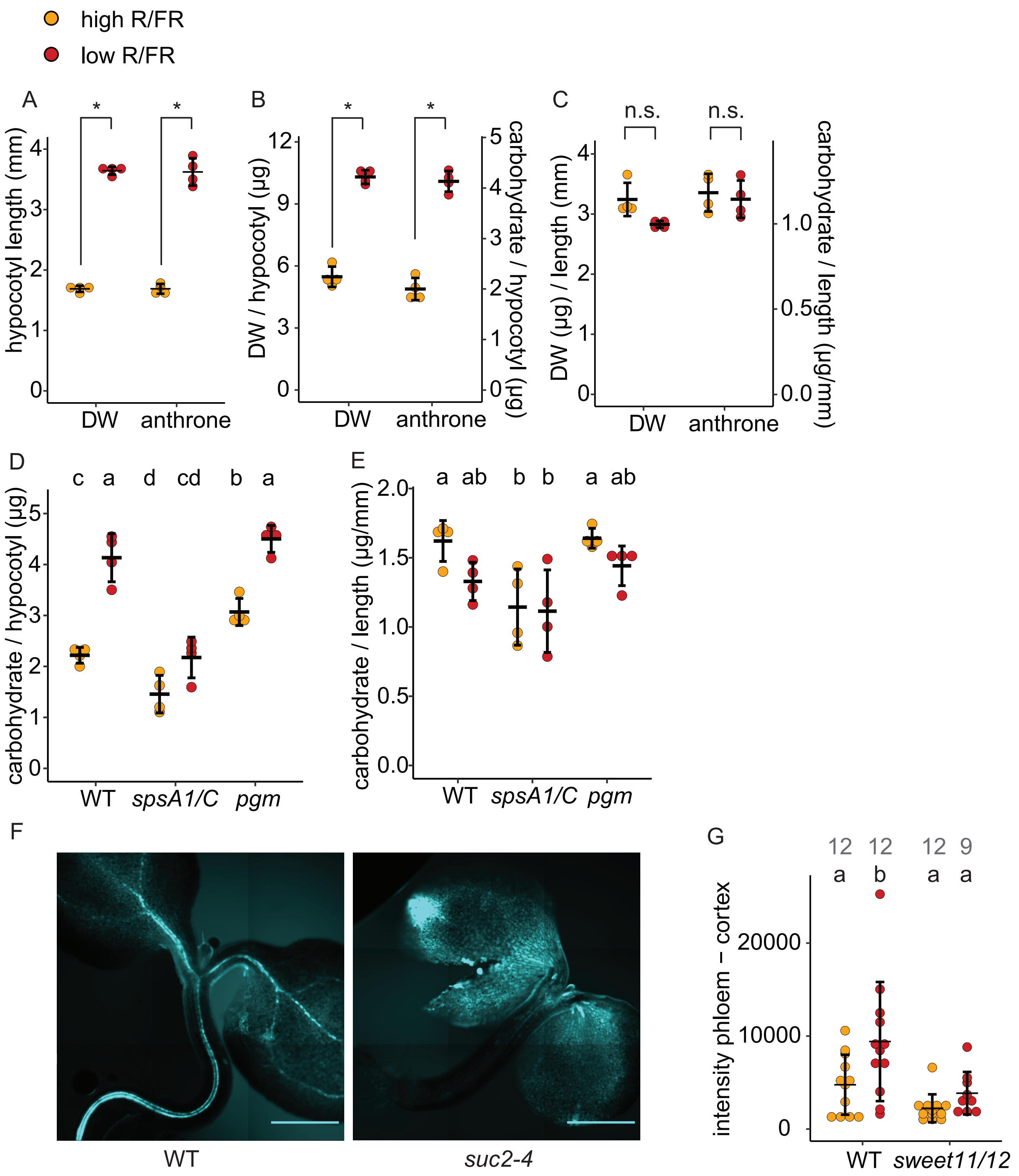


**Figure S2: Neighbor detection is accompanied by enhanced resource allocation to the hypocotyl in Arabidopsis by three independent methods**. A-C): Experimental evidence of hypocotyl biomass increase after 3 days of high vs. low R/FR, using direct DW measurements and carbohydrate content measurement by the anthrone assay. n=4 with 50 to 100 (DW) or 10 to 20 (anthrone assay) hypocotyls per replicate. A) Hypocotyl length of seedlings harvested for both analyses in high (orange) and low (dark red) R/FR. B) DW and carbohydrate content by anthrone assay in high and low R/FR. Carbohydrate content is expressed in µg glucose standard equivalents. C) DW or carbohydrate (anthrone assay) per mm hypocotyl length in high and low R/FR. D) Carbohydrate content of WT, *spsA1/C* and *pgm* hypocotyls by anthrone assay. n=4, with each anthrone assay replicate consisting of 10 to 20 hypocotyls. E) Data from (D) devided by average length of the hypocotyls used. F-G: Measurement of sucrose transport with the phloem mobile dye esculin, which is loaded into the phloem by the SUC2 transporter, therefore the *suc2-4* mutant shows only surface staining, rather than strong fluorescence in the vasculature (F, scale bar: 500µm). G) Quantified fluorescence intensity in the vasculature, with the cortex background subtracted, in high and low R/FR in WT and *sweet11/12*. Grey numbers indicate number of hypocotyls analysed per sample. Statistics in A-C: * $P < 0.05$ (Student's T-test). Statistics in D, E, G: different letters indicate significant difference ($P<0.05$, Tukey's HSD test). Error bars in all plots: standard deviation.
