## Supplementary material for "Internal sugar allocation in response to a shade signal is regulated by concerted action of auxin and sucrose": Figure S3

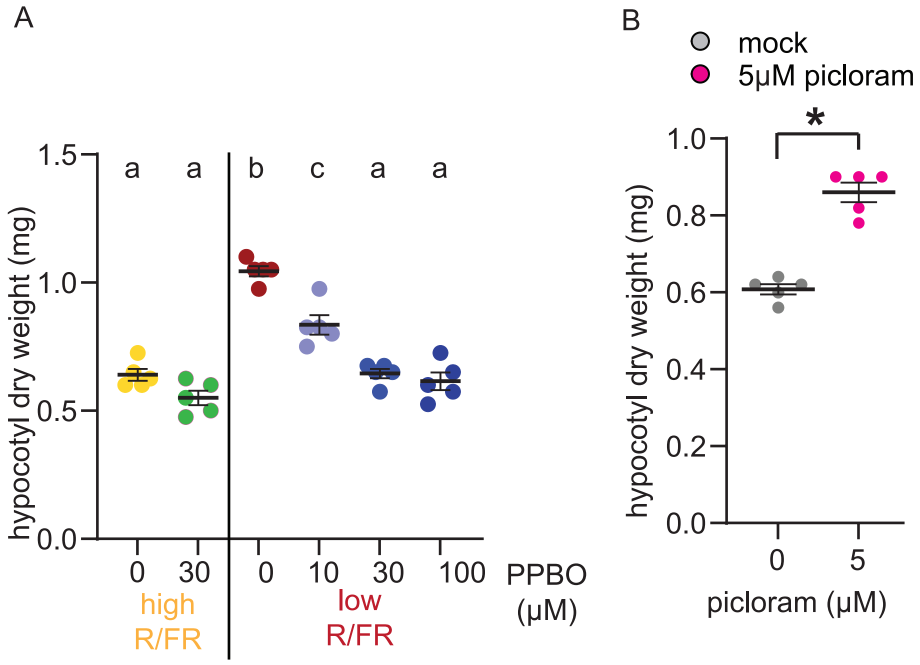


**Figure S3: Auxin is necessary and sufficient for resource allocation to the hypocotyl in *Brassica rapa***. A) Hypocotyl dry weight of 8d old *Brassica rapa* seedlings, grown in high R/FR for 5d followed by 3d in low R/FR with and without PPBO treatment. n=5, where each of the five replicates is the combined dry weight of 4 hypocotyls devided by 4. Different letters indicate statistically significant difference by Tukey HSD-test, perfromed after an ANOVA with significant outcome (P<0.05). B) As in (A), with the last 3d in high R/FR on either mock or picloram supplemented plates. * statistically significant by student's T-test.
