## Supplementary material for "Internal sugar allocation in response to a shade signal is regulated by concerted action of auxin and sucrose": Figure S4

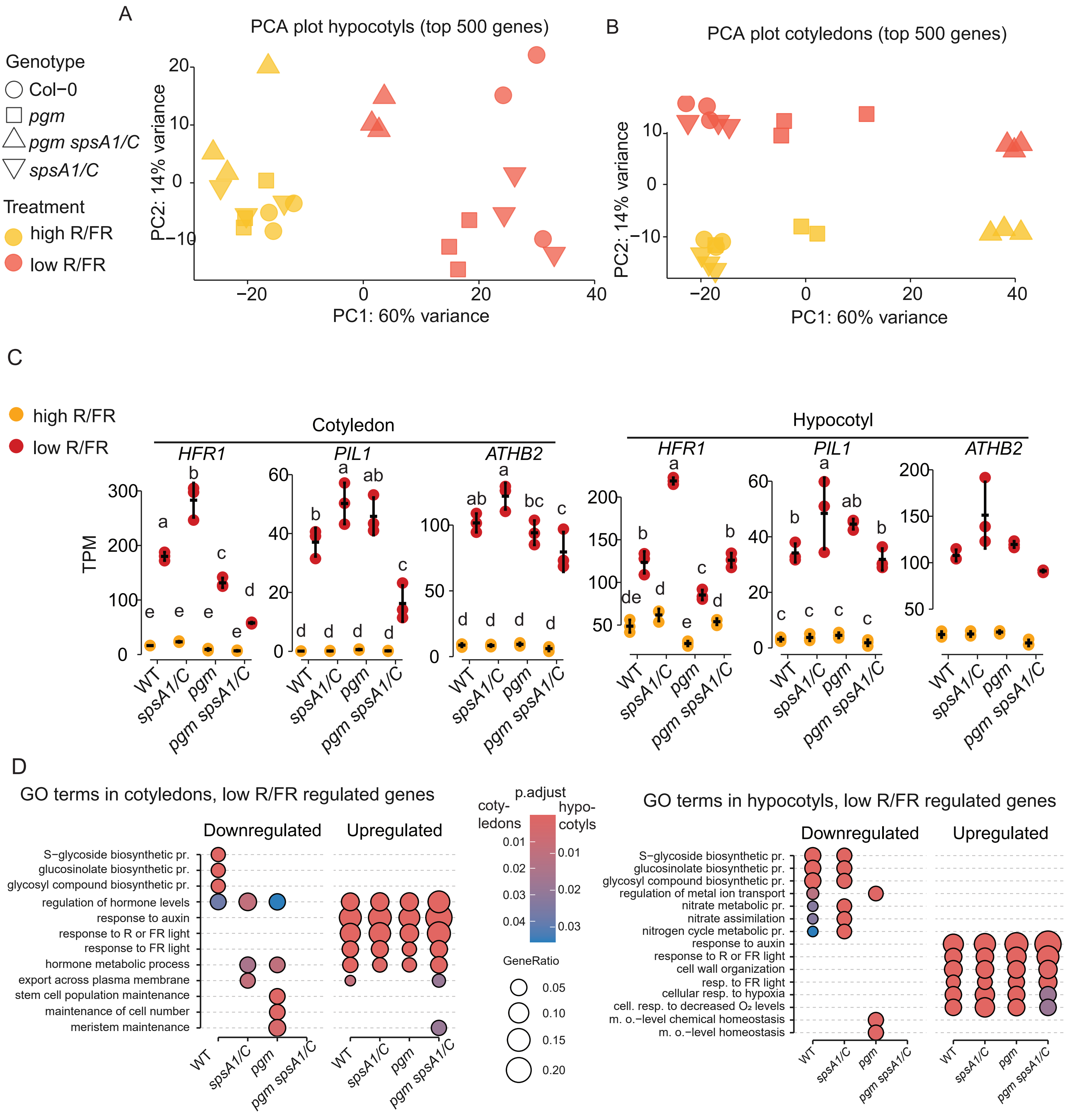


**Figure S4: Additional analysis of RNA-seq dataset**. A-B) PCA of hypocotyl (A) and cotyledon (B) RNA-seq data. C) Expression of the classical neighbor detection-induced shade marker genes *HFR1*, *PIL1* and *ATHB2*. D) Functional enrichment analysis (GO - biological process) of low R/FR responding genes in hypocotyls. pr. = process. m.o. = multicellular organismal. n=3
