## Supplementary material for "Internal sugar allocation in response to a shade signal is regulated by concerted action of auxin and sucrose": Figure S5

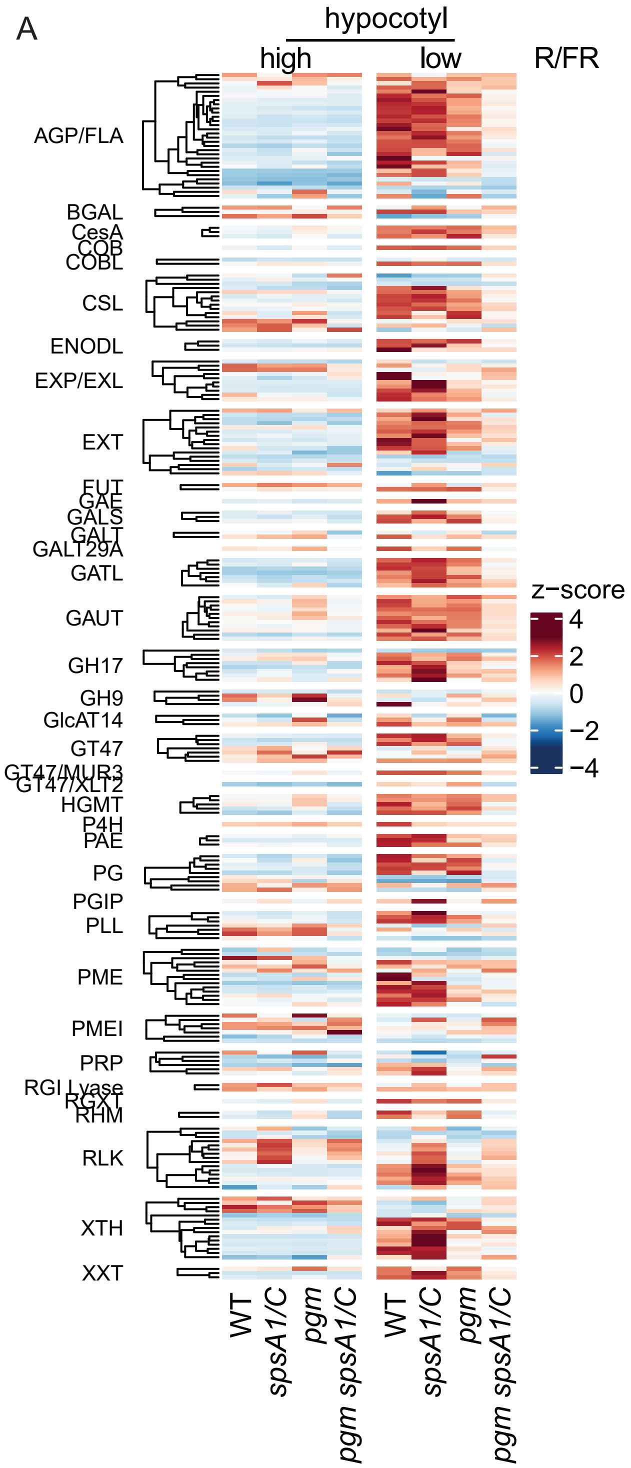


**Figure S5: Expression of cell wall growth related genes.** Heat map of z-scores of cell-wall related differentially expressed genes (Sénéchal *et al.*, 2024).
